# Adeno-associated virus (AAV2) can replicate its DNA by a rolling hairpin or rolling circle mechanism, depending on the helper virus

**DOI:** 10.1101/2023.11.13.566888

**Authors:** Anouk Lkharrazi, Kurt Tobler, Sara Marti, Anna Bratus-Neuenschwander, Bernd Vogt, Cornel Fraefel

**Author notes:** Cornel Fraefel, Winterthurerstrasse 266a, 8057 Zurich, Switzerland. These authors contributed equally to this work (Anouk Lkharrazi, Kurt Tobler).

## Abstract

Adeno-associated virus type 2 (AAV2) is a small, non-pathogenic, helper virus-dependent parvovirus with a single-stranded (ss) DNA genome of approximately 4.7 kb. AAV2 DNA replication requires the presence of a helper virus such as adenovirus type 5 (AdV5) or herpes simplex virus type 1 (HSV-1) and is generally assumed to occur as a strand-displacement rolling hairpin (RHR) mechanism initiated at the AAV2 3’ inverted terminal repeat (ITR). We have recently shown that AAV2 replication supported by HSV-1 leads to the formation of double-stranded head-to-tail concatemers, which provides evidence for a rolling circle replication (RCR) mechanism. We have revisited AAV2 DNA replication and specifically compared the formation of AAV2 replication intermediates in presence of either HSV-1 or AdV5 as the helper virus. The results confirmed that the AAV2 DNA replication mechanism is helper virus-dependent and follows a strand-displacement RHR mechanism when AdV5 is the helper virus and primarily an RCR mechanism when HSV-1 is the helper virus. We also demonstrate that recombination plays a negligible role in AAV2 genome replication. Interestingly, the formation of high molecular weight AAV2 DNA concatemers in presence of HSV-1 as the helper virus was dependent on an intact HSV-1 DNA polymerase.

**Importance:** AAV is a small helper virus-dependent, non-pathogenic parvovirus. The AAV genome replication mechanism was extensively studied in presence of AdV as the helper virus and described to proceed using RHR. Surprisingly, HSV-1 co-infection facilitates RCR of the AAV2 DNA. We directly compared AdV5 and HSV-1 supported AAV2 DNA replication and show that AAV2 can adapt its replication mechanism to the helper virus. Detailed understanding of the AAV replication mechanism expands our knowledge of virus biology and can contribute to increase gene therapy vector production.

## Introduction

Adeno-associated virus (AAV) is a small, non-pathogenic, helper virus-dependent parvovirus (1). The AAV2 genome consists of a 4.7 kb long, single-stranded (ss) linear DNA with two coding regions, *rep* and *cap*. The coding regions are flanked by inverted terminal repeats (ITR) (2). The AAV ITRs consist of 125 nt of palindromic sequences termed A, A’, B, B’, C and C’, followed by a 20 nt unique D sequence adjacent to the terminal resolution site (*trs*) (3). Although the ITRs are non-coding, they constitute an important part of the AAV genome, as they self-assemble into a double-hairpin structure that serves as a primer for DNA replication and as a packaging signal (4,5). Although AAV can infect host cells in the absence of helper factors, it can only replicate if the cellular environment undergoes a dramatic change (6,7) or in presence of helper factors provided by specific helper viruses. These helper viruses include adenoviruses (AdV) and herpesviruses (HSV) (8–10) amongst others. Additionally, cellular stress, such as UV radiation or carcinogens can induce AAV2 replication to a certain extent (7,11).

HSV-1 is a large, enveloped virus of approx. 200 nm in diameter with a linear 152 kbp long dsDNA genome (12). The genome is replicated by the HSV-1 polymerase UL30/UL42 and packaged into preformed capsids. Interestingly, viral genomes are only packaged if longer-than-unit-length viral DNA is present (13), therefore concatemerization of HSV-1 genomes must take place before packaging. In early work it was suggested that the HSV-1 genome circularizes and enables rolling circle replication (RCR) which inherently leads to concatemers (14). More recent work suggests concatemerization through recombination mediated by ICP8 and UL12 (15,16).

The minimal helper factors necessary to support AAV replication in AAV and HSV-1 co-infected cells include the HSV-1 helicase primase complex composed of the UL5, UL8, and UL52 proteins and the HSV-1 ssDNA binding protein ICP8 (17). Although the HSV-1 polymerase (HSV-1 pol), consisting of UL30 and UL42, is not strictly necessary for AAV genome replication it was shown to enhance the process (18,19).

AdVs are nonenveloped viruses with a linear, non-segmented dsDNA genome of about 35-36 kbp (20). The AdV genome is replicated by the AdV polymerase through a protein-primed strand-displacement mechanism (reviewed in (21)). The essential helper factors provided by AdV for AAV genome replication include E1A and E2A which are needed for activation of the AAV *p5* and *p19* promoters (22–24), E1B55K which in concert with E4orf6 promotes second strand synthesis and viral DNA replication (25,26) and the VA RNA which is required for efficient synthesis of AAV structural proteins (27). Interestingly, in AdV supported AAV genome replication, the cellular polymerase δ was shown to be essential, while the AdV polymerase was not (28,29).

AdV supported AAV genome replication is suggested to occur in a rolling hairpin mechanism (RHR) using the ITR as primer to initiate second-strand synthesis, which leads to the formation of a covalently closed duplex structure (30). Nicking by Rep68 or Rep78 at the *trs* produces a 3’-end that serves as primer for synthesis of a new ITR, resulting in the resolution of the duplex structure (31). In a process termed re-initiation, hairpins form at the ends of the genome and present a new 3’-end primer for DNA synthesis that displaces and releases a single stranded AAV genome. The newly synthesized AAV DNA is then packaged into preformed capsids (32). There is ample experimental evidence supporting this model. For example, two distinct ITR orientations, termed “flip” and “flop”, have been observed, with their relative positioning found to be independent of each other (2,33). The ITRs can act as origin of DNA replication (3,5,34). Covalently closed double-stranded (ds) monomer structures as predicted by the RHR model have indeed been isolated from AAV-infected cells (35) and site-specific nicking of the *trs* has been observed (31,36,37).

Recently we provided evidence that AAV2 replicates its DNA using a RCR rather than a RHR mechanism when HSV-1 is the helper virus (38), suggesting that AAV2 can adapt the genome replication strategy to the helper virus present. Here, we directly compared the AAV2 DNA replication intermediates formed in presence of either HSV-1 or AdV5. The data is consistent with a preferential RHR in presence of AdV5 and preferential RCR in presence of HSV-1. We also demonstrate that recombination plays a negligible role in AAV2 genome replication.

## Results

### Nanopore sequencing reveals different AAV genome replication intermediates depending on the helper virus, HSV-1 or AdV5

We have recently shown that HSV-1 co-infection facilitates RCR of AAV2 DNA (38), while earlier reports have described AAV to replicate via a RHR mechanism. To investigate this discrepancy, we directly compared AAV2 genome replication when supported by either HSV-1 or AdV5. To analyze AAV2 DNA replication intermediates, we employed Oxford Nanopore sequencing, following the previously described method by Meier et al. (38). Specifically, cells were co-infected with AAV2 (MOI 100) and either HSV-1 (MOI 0.1) or AdV5 (MOI 0.1). After 48 h, the cells were harvested, and extrachromosomal DNA was extracted using the Hirt protocol (39). To minimize background of input AAV2 DNA the MOI was kept low. Sequencing results showed different DNA replication intermediates, that were classified according to the previously described categories 1) monomers, 2) dimers, 3) head-to-tail (HT) repeats, 4) alternating repeats, 5) HT and alternating repeats, 6) ITR repeats and 7) others, that could not be classified in one of the categories 1-6 (38). HT repeats are indicative of RCR, while alternating repeats may arise from RHR in absence of terminal resolution. The results are shown in Table 1 and can be summarized as follows: the majority of reads consisted of AAV2 genome monomers in both HSV-1 (33%) and AdV5 (29%) supported replication. There was an approx. five-fold higher percentage of HT repeats (11%) compared to alternating repeats (2%) in HSV-1 co-infected cells. Cells co-infected with AdV5 showed more alternating repeats (2%) than HT repeats (1%). Interestingly, AdV5 co-infection induced the generation of ITR repeats much more efficiently (29.6%) than HSV-1 co-infection (2.4%).

**Table 1.** Read analysis of sequencing data from cells co-infected with AAV2 (MOI 100) and either HSV-1 (MOI 0.5) or AdV5 (MOI 0.5) at 48 hpi. Table shows ratios of the different categories of AAV2 DNA replication products of a total of 500 reads per sample.

| Category | 1 | 2 | 3 | 4 | 5 | 6 | 7 |
| --- | --- | --- | --- | --- | --- | --- | --- |
|  | Monomer | Duplex | Head-to-tail Repeats | Alternating Repeats | Head-to-tail and Alternating Repeats | ITR Repeats | Others |
| Sample | ratio | ratio | ratio | ratio | ratio | ratio | ratio |
| AAV2/HSV-1 | 0.330 | 0.256 | 0.110 | 0.024 | 0.020 | 0.024 | 0.236 |
| AAV2/AdV | 0.290 | 0.130 | 0.010 | 0.020 | 0.004 | 0.296 | 0.250 |

**Table 2.** Primer sequences used for Southern blot probe synthesis. Table indicates generated probe length and hybridiza?on temperature for each probe.

| Probe | Forward primer | Reverse primer | Probe length | Hybridization temperature |
| --- | --- | --- | --- | --- |
| *rep* | 5'-gaa cgc gat<br>atc gca gcc gcc<br>atg ccg gg-3' | 5'-gga tcc gaa<br>ttc act gct tct ccg<br>agg taa tc-3' | 735 bp | 48.9°C |
| CFP/GFP | 5'-ctg acc ctg<br>aag ttc atc tgc-3' | 5'-gga tct tga<br>agt tgg cct tg-3' | 378 bp | 50.3°C |
| mCherry | 5'-agg acg gcg<br>agt tca tct ac-3' | 5'-ttc cac gat ggt<br>gta gtc ctc-3' | 295 bp | 51.1°C |

### Southern analysis confirms differential AAV2 DNA replication mechanisms depending on the helper virus

AAV2 replication intermediates were also investigated by Southern blot of Hirt DNA extracted from cells co-infected with AAV2 and either HSV-1 or AdV5. Some of the samples were treated with *Hin*dIII to investigate the nature of the concatemeric replication intermediates. *Hin*dIII cuts the ds AAV2 genome within the *rep*-sequence, specifically at nucleotide position 1882 (NC_001401, Fig. 1A), and yields the following predicted DNA fragments: 1.9 kb and 2.8 kb for monomers, 3.8 kb and 5.6 kb for head-to-head (HH) concatemers, and 4.7 kb for HT concatemers. Because of the specificity of the hybridization probe (Fig. 1A) only the bands at 1.9 kb, 3.8 kb and 4.7 kb can be detected. In both untreated samples, bands at approx. 3 kb, which represent the ss AAV2 genome, and at approx. 4.7 kb and 10 kb, which correspond to ds monomeric and dimeric AAV2 DNA, respectively, were observed (Fig. 1B). Higher molecular weight bands representing AAV2 genome concatemers were observed only in presence of HSV-1 as the helper virus. *Hin*dIII treatment resulted in the disappearance of all bands representing ds dimers and higher order multimers of the AAV2 genome. Instead, a strong band at 4.7 kb, which is expected from cleavage of HT-linked concatemers that arise from RCR, appeared in the Hirt DNA prepared from AAV2 and HSV-1 co-infected cells. Additional bands appeared at 3.8 kb and 1.9 kb which are consistent with cleavage of HH-linked concatemers arising from RHR and of ds AAV2 genome monomers, respectively. In the samples from AAV2 and AdV5 co-infected cells, *Hin*dIII cleavage produced strong bands at 3.8 kb, which is expected from cleavage of HH-linked concatemers, and at 1.9 kb, which is expected from cleavage of ds AAV2 monomers. A much weaker band at 4.7 kb may also represent a cleavage product from HT-linked concatemers. Overall, the data corroborates the sequencing data and indicates that AAV2 replicates predominantly using the RCR mechanism in presence of HSV-1 and the RHR mechanism in presence of AdV5.

**Fig. 1.**
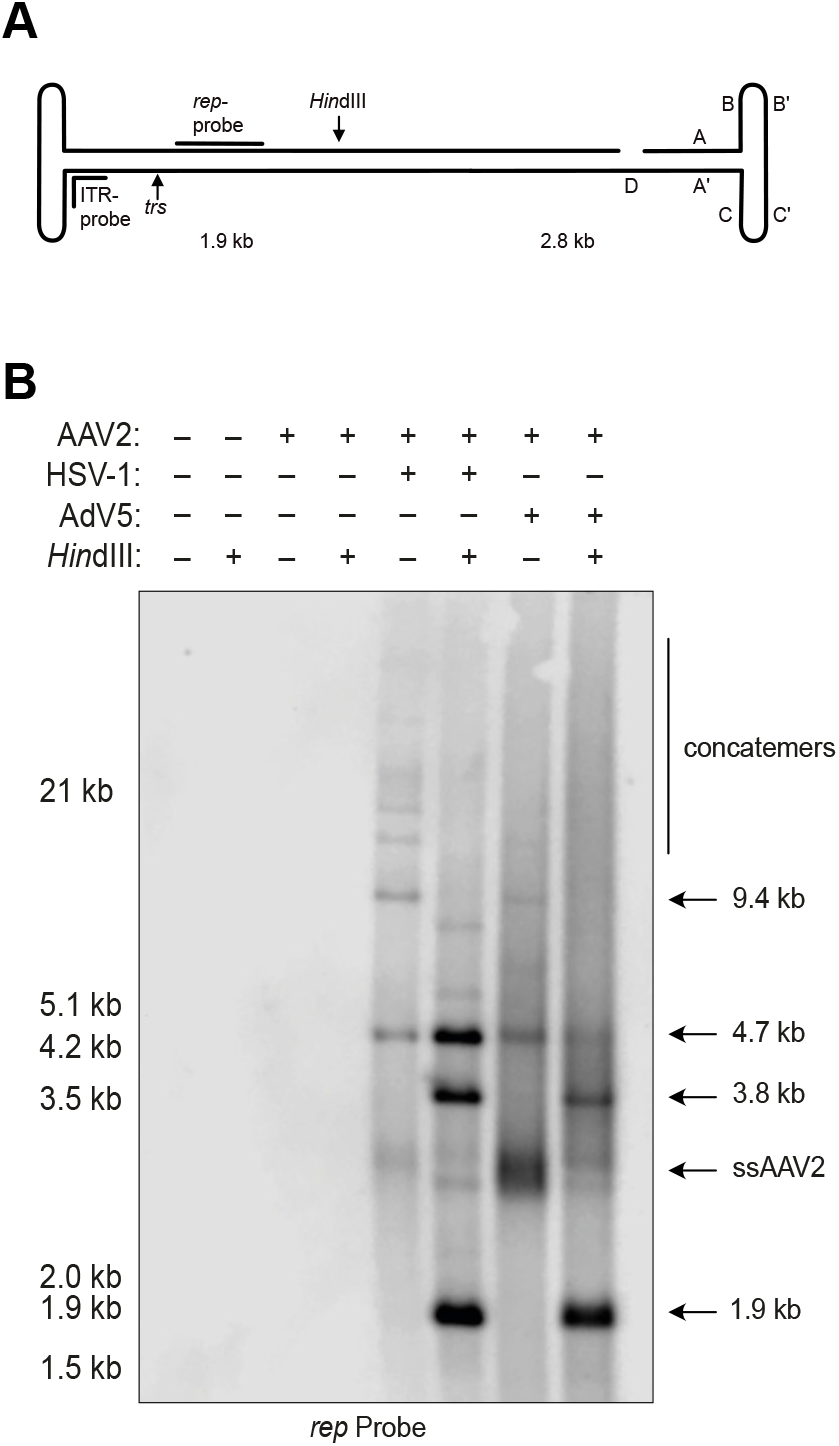
Southern blot of AAV2 DNA replication intermediates. (A) Schematic representation of the AAV2 genome with ITR regions, *trs*, *Hin*dIII restriction site and probe binding sites indicated. (B) Southern blot of untreated or *Hin*dIII treated Hirt DNA extracted from cells infected with AAV2 (MOI 100) or co-infected with AAV2 (MOI 100) and either HSV-1 (MOI 0.5) or AdV5 (MOI 0.5) at 48 hpi. Bands were visualized with a *rep*-specific probe.

### AdV5 co-infection leads to the generation of vast amounts of ITR repeats

To exclude the possibility that the ITR repeats identified by Oxford Nanopore sequencing (Table 1) are due to a sequencing artifact, we designed a hybridization probe specific for the C and A sequence within the ITR (Fig. 2A). Southern analysis showed robust staining of low molecular weight bands in samples from AAV2 and AdV5 co-infected cells. These bands were observed also in samples from AAV2 and HSV-1 co-infection but with a much lower intensity. Distinct further bands at approx. 3 kb and 4.7 kb were detected in both co-infections and may represent ss and ds AAV2 DNA monomers, respectively. An additional band of approx. 10 kb was observed in the sample from AAV2 and HSV-1 co-infected cells only and may represent the ds dimer of the AAV2 DNA (Fig. 2A).

**Fig. 2.**
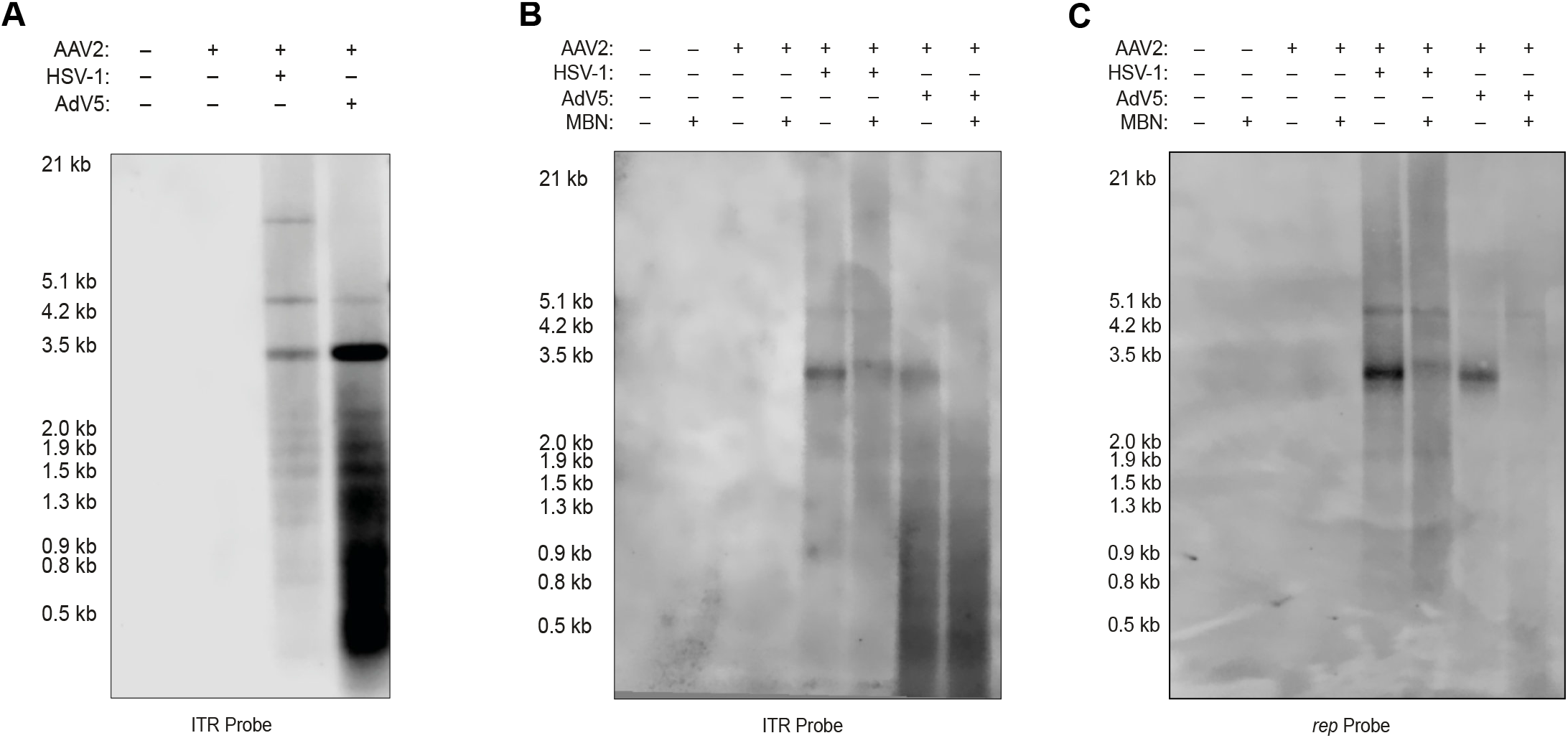
Southern analysis of ITR repeats in total cells or supernatant. Southern blot of Hirt DNA extracted from (A) total cells or (B and C) supernatant infected with only AAV2 (MOI 100) or co-infected with AAV2 (MOI 100) and HSV-1 (MOI 0.5) or AdV5 (MOI 0.5) at 48 hpi. Bands were visualized with either (A and B) an ITR specific probe or (C) a *rep* specific probe.

Interestingly, using the ITR-probe (Fig. 2B) but not the *rep* probe (Fig. 2C), strong staining of sub genome-length low molecular weight, Mung bean nuclease resistant DNA was observed when DNA was extracted from progeny AAV2 particles purified from AAV2 and AdV5 co-infected cells.

### ITRs in head-to-tail concatemers show the same orientation

To further investigate, whether HT concatemers of the AAV2 genome are a product of RCR, the nucleotide sequence was examined in greater detail. Hirt DNA from cells infected as described above was analysed using PacBio sequencing, as this method offers higher accuracy on single base level compared to Oxford Nanopore sequencing (40,41). The sequencing data revealed that the ITR regions joining two AAV2 genomes in the concatemeric DNA (HT and alternating repeats) consisted of a modified ITR (TRT) containing an additional flanking D sequence (Fig. 3B and C), which has been previously described in circular AAV genomes isolated from infected cells (3). Further, multiple sequence alignment (MSA) of six HT reads (Fig. S1A) show that the ITRs occur indeed exclusively in either a flip or a flop orientation in one read. This further supports RCR as mechanism for HT concatemers, as the same orientation of the TRT would be expected from RCR using cAAV genomes as substrate. The ITRs in alternating repeats (Fig. S1B) showed a random distribution of orientation (Fig. 3C). This finding is consistent with RHR in absence of *trs* resolution, as within one AAV genome the orientation of the ITRs is random and independent of each other (33). Interestingly, one read with alternating repeats was found, showing an additional CC’ sequence in two of the five ITR regions within the read (Fig. 3C).

**Fig. 3.**
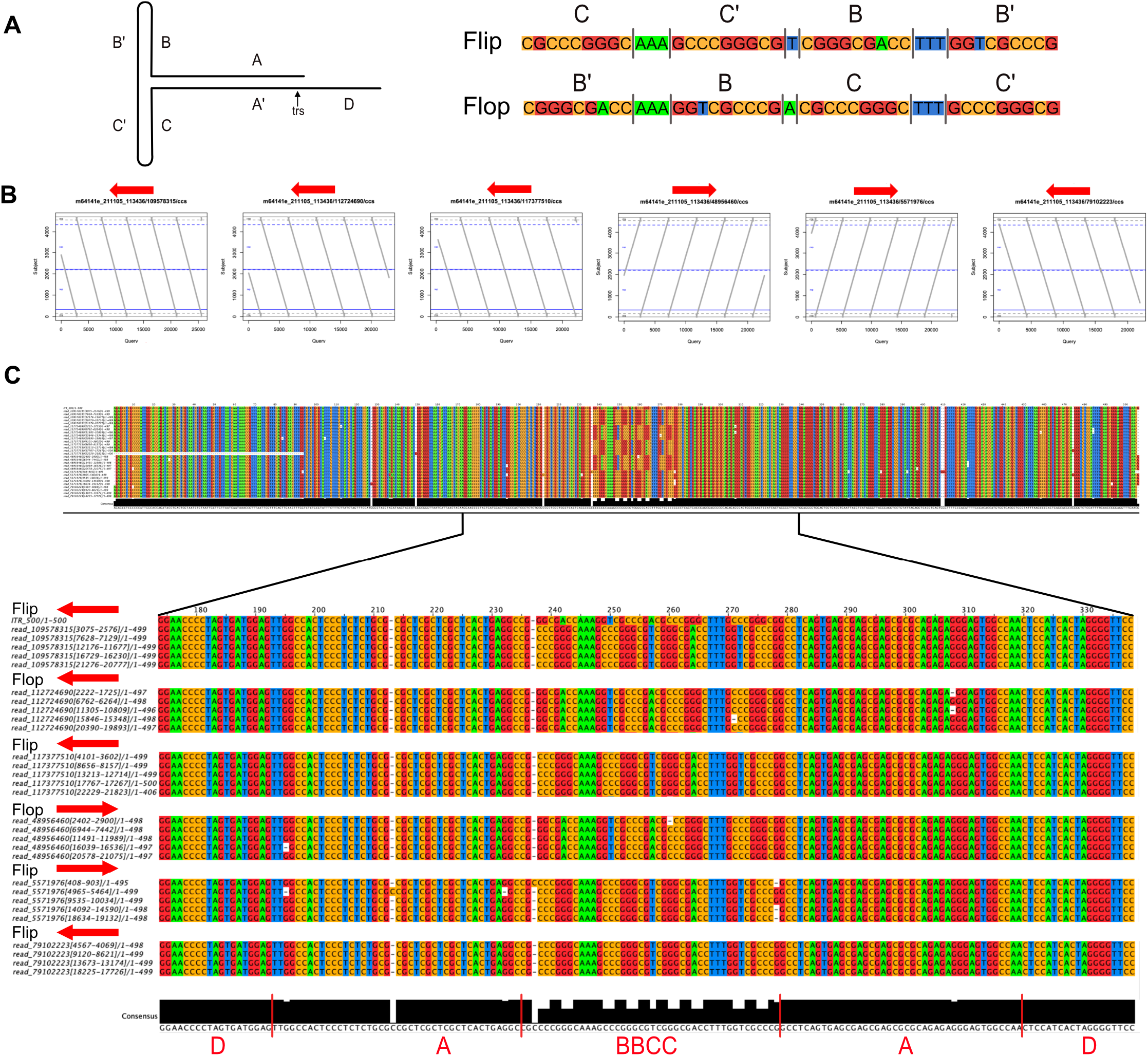
MSA analysis of ITR regions. (A) Schematic representation of the Flip and Flop orientation of the BBCC region of the ITR. Colors represent different bases: C, orange; G, red; A, green; T, blue. The ITR regions of the six longest (B) HT repeats and (C) alternating repeats from HSV-1 co-infection were aligned and the orientations indicated. Each block represents one read.

### The contribution of recombination to the formation of AAV2 genome concatemers is negligible

AAV2 genome concatemers could arguably also arise from genome recombination events. However, the probability of recombination leading to the formation of long, HT concatemers of the AAV2 DNA would be exceedingly low, as previously discussed by Meier et al. (38). Nevertheless, to show if and to what extent recombination plays a role in HSV-1 supported AAV2 genome replication, Hirt DNA of cells infected with two different recombinant AAV vectors (rAAV), encoding for either GFP (rAAVGFP) or mCherry (rAAVmC) and co-infected with a recombinant HSV-1 encoding Rep and Cap (rHSV-1RC, Fig. S2) to support rAAV replication were analyzed by Southern blot (Fig. 4, A-C). Four regions with lengths ranging from 120 bp to 700 bp are virtually identical between the two different rAAV vector genomes and would readily support homologous recombination. To reveal the nature of the concatemers some samples were treated with *Sal*I which cuts the ds rAAV genomes at nucleotide position 815 (Fig. 4A) and yields the following predicted DNA fragments: 0.8 kb and 1.2 kb from monomers of rAAVGFP or 0.8 kb and 3 kb from monomers of rAAVmC, of which only the bands at 1.2 kb and 3 kb can be detected because of the specificity of the hybridization probe. *Sal*I digestion of HT concatemers of rAAVGFP and rAAVmC genomes would yield detectable bands at 2 kb and 3.8 kb, respectively. *Sal*I digestion of HH concatemers of rAAVGFP and rAAVmC genomes would yield detectable bands at 2.4 kb and 6 kb, respectively. Cleavage of a recombined, mixed HH concatemer, would yield a band at 4.2 kb. Cleavage of a recombined, mixed HT concatemer would yield bands at 2 kb or 3.8 kb, but these could not be differentiated from *Sal*I digested HT concatemers of rAAVmC or rAAVGFP. Figure 4, B and C, shows that co-infections with each rAAV individually or in combination and the helper virus yielded the expected patterns for the respective vector genomes, including high molecular weight concatemers. S*al*I treatment resulted in the disappearance of all bands representing ds dimers and higher order multimers of the rAAV genomes. Instead, strong bands at 2 kb (Fig. 4B) and 3.8 kb (Fig. 4C), which are expected from cleavage of RCR products, appeared. Additional bands appeared at 2.4 kb (Fig. 4B) and 6 kb (Fig. 4C), which are consistent with RHR products. Importantly, at 4.2 kb, the expected size for a recombined HH fragment, no band was detected with either probe.

**Fig. 4.**
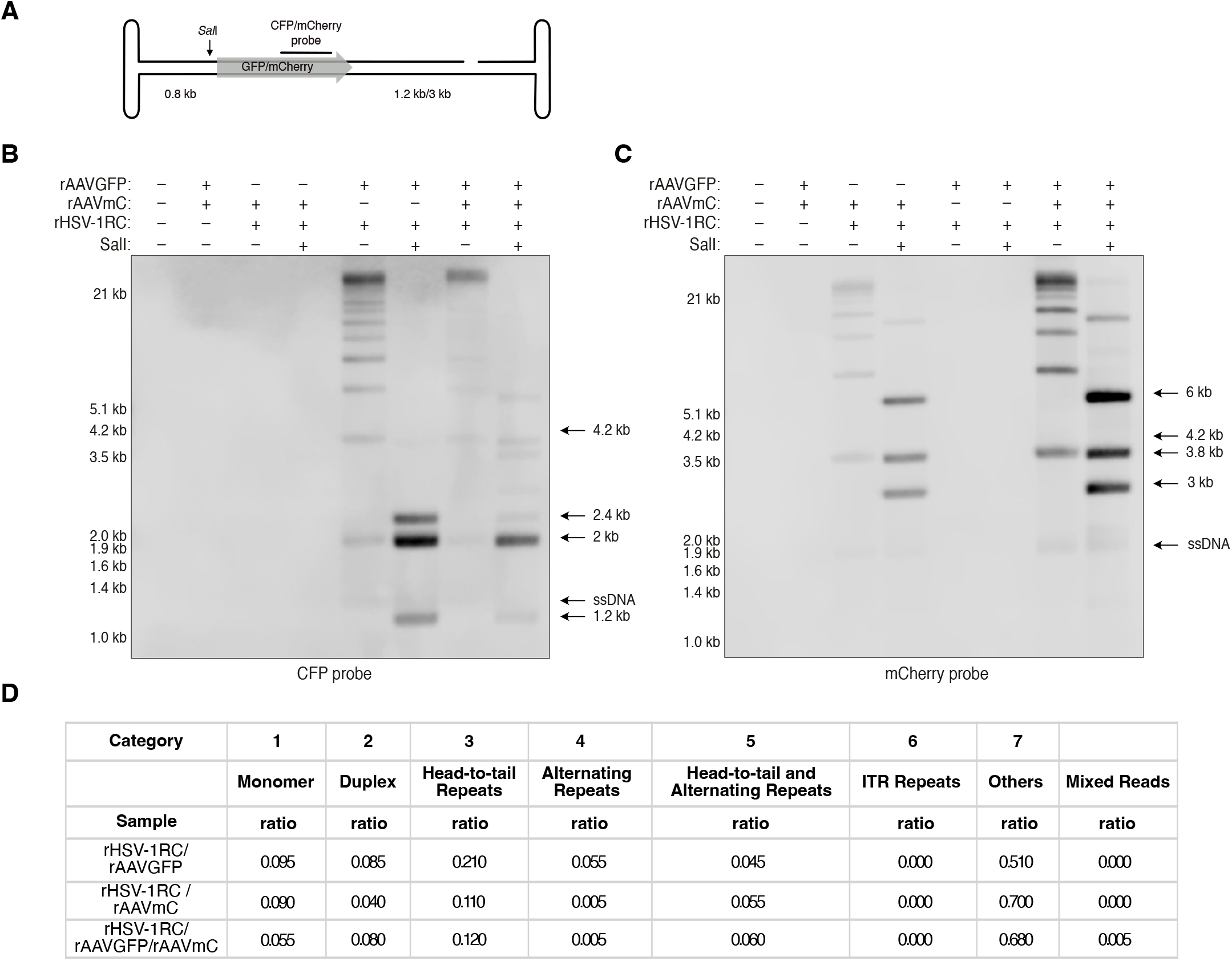
Southern blot and nanopore read analysis of recombination efficiency. (A) Schematic representation of the rAAV genome with *Sal*I cut site, expected fragment sizes, and probe binding site indicated. Transgenes are indicated as grey arrow. Southern blot of untreated or *Sal*I treated Hirt DNA from cells infected with rAAVGFP (MOI 100, 2 kb), rAAVmC (MOI 100, 3.8 kb) and rHSV-1RC (MOI 0.5) in indicated combinations at 48 hpi. Sizes of HH and HT concatemers or monomer fragments in *Sal*I treated samples are indicated with arrows. The arrow at 4.2 kb indicates the predicted size of a *Sal*I digested mixed fragment. AAV2 DNA was visualized with either (B) a CFP probe or (C) a mCherry probe. (D) Read analysis of sequencing data of cells infected as described above. Table shows ratio of the different categories of a total of 200 reads.

As recombination levels might be below the detection limit of the Southern blot, we additionally investigated recombination on a single genome level using Oxford Nanopore sequencing. Cells were infected as described above and Hirt DNA was prepared and sequenced. Similar to the co-infections with wild-type (wt) AAV2, in all co-infections with rAAVs more HT repeats (21%, 11% and 12%) were present compared to alternating repeats (5.5%, 0.5%, 0.5%) (Fig. 4D and S3), indicating that replication preferentially occurs by RCR. The mixed reads in cells co-infected with rAAVGFP, rAAVmC and HSV-1RC showed a ratio 0.5% indicating that recombination plays a minimal role in concatemer formation. Interestingly, when using rAAV instead of wt AAV2 in co-infections with a helper virus, most reads belonged to category 7 and no ITR repeats were observed (Fig. 4D).

### The HSV-1 polymerase is essential in AAV2 genome concatemer formation

Depending on the helper virus supporting AAV replication, different cellular and viral proteins are recruited to AAV2 replication compartments (42,43). The HSV-1 polymerase is not strictly necessary for AAV replication but was demonstrated to strongly enhance it (18,19,44). As HSV-1 replicates its DNA using a RCR mechanism and requires HT concatemers for packaging (13), we investigated the possibility that the HSV-1 polymerase is involved in the formation of HT repeats of the AAV2 genome. For this, cells were co-infected with either wt HSV-1 or an HSV-1 mutant (HSVΔUL30) in which the catalytic subunit (UL30) of the HSV-1 polymerase complex is non-functional. In a second step, cells infected with HSVΔUL30 were reconstituted by transfection of an UL30 encoding plasmid. The results of Southern analysis and Nanopore sequencing indeed imply an important role for the HSV-1 polymerase in concatemer formation, as HT concatemers of the AAV2 genome were not found in presence of HSVΔUL30 as the helper virus (Fig. 5A and B) but were detected after reconstitution of UL30 (Fig. 5A).

**Fig. 5.**
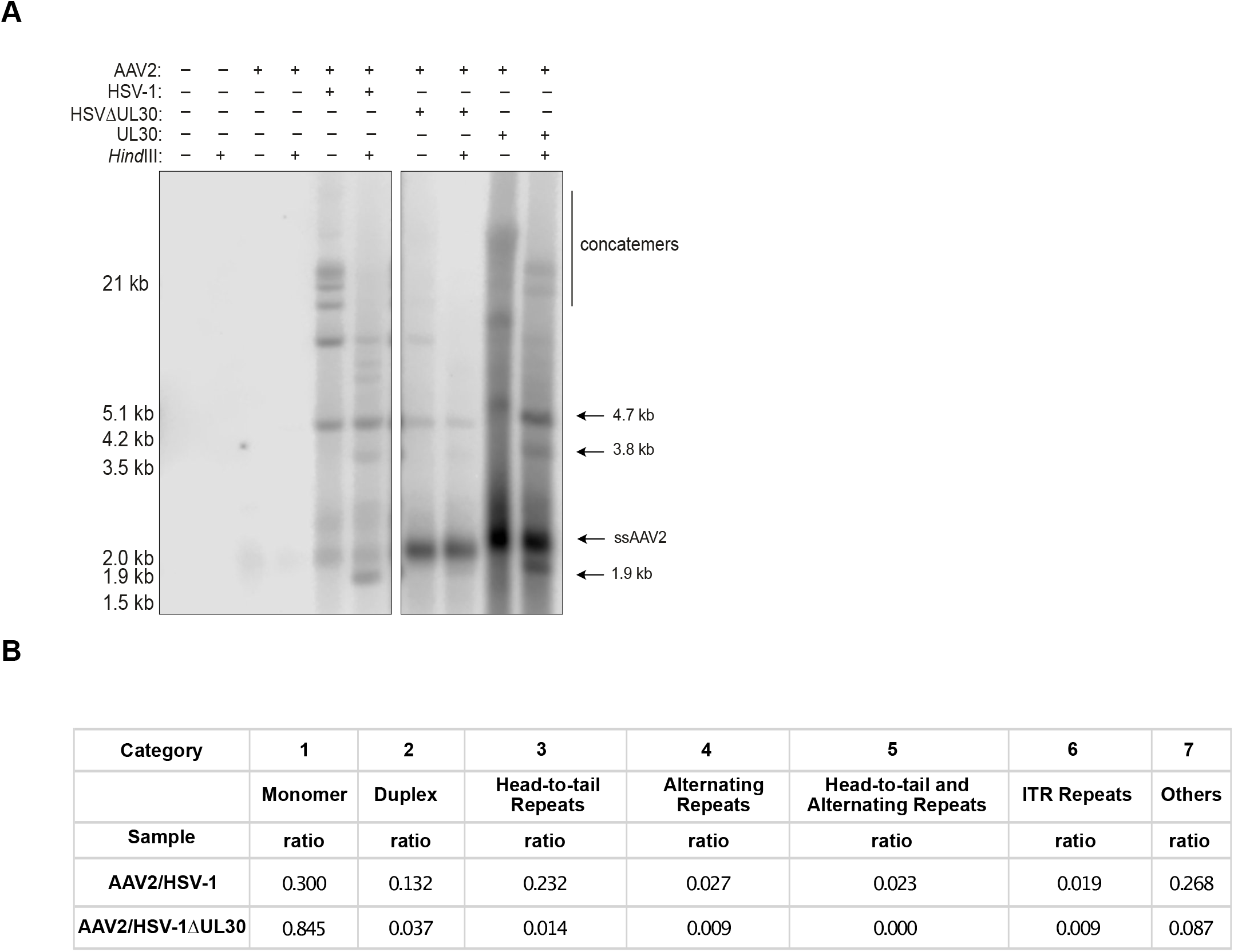
Southern blot and Oxford Nanopore read analysis of AAV2 DNA replication intermediates. (A) Southern blot of untreated or *Hin*dIII treated Hirt DNA from cells infected with AAV2 or co-infected with AAV2 (MOI 100) and HSV-1 (MOI 0.1), HSV-1ΔUL30 (MOI 1), or HSV-1ΔUL30 (MOI 1) with reconstituted UL30. (B) Sequencing read analysis of Hirt DNA from cells co-infected with AAV2 and HSV-1 or HSV-1ΔUL30. Table shows ratio of the different categories of a total of 200 reads.

## Discussion

In presence of AdV as the helper virus, AAV DNA replication is assumed to proceed in a RHR mechanism (30,35,45,46). However, AAV may adapt its replication process depending on the cellular environment and the helper virus, as we recently showed that HSV-1 co-infection facilitates RCR of the AAV2 genome (38). Here we directly compared the formation of AAV2 DNA replication intermediates in presence of the two different helper viruses and analyzed the junctions between concatemeric genomes in detail. The sequencing data as well as the results from Southern blotting revealed a much higher ratio of HT concatemers, the products of RCR, compared to alternating repeats, the products of RHR, in HSV-1 co-infection while the opposite was true in AdV5 co-infection.

The formation of divergent sets of AAV2 DNA replication products is likely due to the different helper functions provided. The essential helper factors from AdV include E1a (22), E1b55k (25), E2a (47), E4orf6 (48) and the VA RNA (27); the essential HSV-1 helper factors consist of the helicase-primase (HP) complex, composed of UL5/UL8/UL52, and the ssDNA binding protein ICP8 (17). Interestingly, neither the AdV DNA polymerase nor the HSV-1 DNA polymerase are required for AAV genome amplification, but the HSV-1 DNA polymerase was shown to strongly enhance AAV genome replication (18,19,44). The HSV-1 DNA polymerase together with ICP8 and the HP complex can confer RCR to circular plasmids even in absence of an HSV-1 origin of DNA replication or the HSV-1 origin binding protein (49). The AAV DNA has indeed been shown to readily circularize within the host cell (50–52) thereby provides a potential template for RCR. Moreover, the circular AAV genomes (cAAV) contain a TRT which constitutes a functional origin for AAV DNA replication (3). Our sequencing data revealed that the AAV2 genomes in the HT concatemer are not only joined by TRTs, but also that all TRTs in one molecule have the same orientation, either all flip or all flop. These two observations concerning the TRTs further support that AAV2 DNA synthesis proceeds as a RCR mechanism, at least in presence of HSV-1, and suggest that cAAV is the most likely substrate. High molecular weight AAV2 genome concatemers were only detected in HSV-1 supported AAV2 genome replication but not in AdV5 supported replication. This might be possibly due to AdV5 proteins that are able to counteract concatemer formation. For example, AdV E4orf6 has been shown to stabilize the AdV genome and prevent unwanted concatemerization. During AdV infection, E4orf6 in complex with E1B55k inhibits p53-mediated anti-viral responses and non-homologous end joining (NHEJ)-dependent processes of concatemerization of the AdV genome through targeted degradation of p53 and meiotic recombination 11 (MRE11) (53,54). As an essential helper factor in AAV genome replication (48,55), AdV E4orf6 may have a similar role in the inhibition of AAV2 genome concatemerization. However, our finding that with either helper virus the contribution of recombination to the formation of AAV genome concatemers is negligible argues against this possibility as E4orf6 blocks concatemer formation via blocking NHEJ-mediated recombination (53,54).

Alternatively, lack of HT concatemers of the AAV2 genome in presence of AdV5 might not be due to a different replication mechanism but rather due to more efficient terminal resolution, which prevents concatemer formation.

The question remains whether HT concatemers can yield single stranded, packageable, and infectious AAV genomes and/or contribute to AAV infection as templates for viral gene transcription. Packaging into AAV capsids would require the excision from a concatemer of a single-stranded monomeric AAV2 genome containing a complete ITR at each end. A mechanism that yields such genomes from concatemers is indeed conceivable if considering that cAAV containing a single TRT represents the simplest form of a HT concatemer. Musatov et al. (56) demonstrated that cAAV can give rise to packageable AAV genomes and proposed the following model for this: Rep68/78 can bind to the A sequence and initiate RCR by nicking at the *trs*. This is followed by an extension of the newly generated 3’ end and simultaneous strand displacement. After the replication fork has completed a full circle, the displaced strand is cleaved by Rep, resulting in a single-stranded monomer with a complete ITR on the 5’-end and a 3’-end with only a D sequence. The incomplete ITR can be repaired via a single-stranded panhandle intermediate formed by annealing of two inverted D sequences and extension of the incomplete strand, a mechanism that has indeed been described for AAV (5).

There is a striking difference in the abundance of AAV2 ITR repeats formed in presence of HSV-1 and AdV5. Interestingly, the ITR repeats were also found in AAV2 progeny, encapsidated or bound to the outside of the capsid, and may therefore play a role early in AAV2 infection. For example, AAV ITRs can activate the DNA damage response (DDR) involving Ataxia-Telangiectasia Mutated (ATM) and RAD3-related (ATR) kinases, that inhibit cell cycle progression and lead to apoptosis in a p53 dependent manner (57,58). Extra-viral DNA has been shown to induce Toll-like receptor 9 (TLR9)-dependent immune responses in human plasmacytoid dendritic cells, which was inhibited by DNase treatment of the AAV stocks (59).

Proteins of the DDR may be involved in the formation of ITR repeats as DNA-dependent protein kinase catalytic subunit (DNA-PKcs) and Artemis-associated endonuclease (Artemis) have been shown to cleave ITR hairpins of rAAV vectors in a tissue-dependent manner (60). Besides DNA-PKcs, Artemis can also be activated through ATM (61). The resulting structure can be cleaved at the *trs* by Rep68/78 and the newly generated 3’-end can be extended, resulting in a fragment containing two ITRs. Subsequently the ITRs can reanneal, forming a double hairpin structure and thereby provide self-priming activity. The 3’-end can then again be extended. Following this mechanism, it is conceivable that the ITRs cleaved by DNA-PKcs and Artemis can serve as substrate for the observed ITR repeats. The difference in ITR repeat ratios in AdV5 versus HSV-1 supported AAV2 genome replication may be due to differences in DNA-PKcs levels, as this kinase was found to be the primary mediator of damage signaling in response to AAV genome replication in AdV co-infection (62). In HSV-1 co-infection DNA-PKcs was shown to be degraded in an ICP0 dependent manner (63,64), although at a delayed rate in co-infection with AAV2 (65). This delay may be important for HSV-1 supported RCR of the AAV genome, as AAV genome circularization, a pre-requisite for RCR (66), was shown to be impaired in DNA-PKcs deficient SCID mice (67–69). Furthermore, the ability to induce a significant DDR was linked to the p5 region on rAAV2 vectors as vectors lacking the p5 region of AAV2 were not capable of inducing such a response (70). Indeed, in infections with rAAV2s, that do not contain the p5 region, we did not find ITR repeats, further linking the formation of ITR repeats to the DDR.

In this study we provide evidence that AAV2 can adapt its replication mechanism depending on the helper virus. Several differences in the ratios of specific replication intermediates such as concatemers and ITR repeats were found. The biological relevance of these need further investigations. Besides different helper factors, differences in Rep levels/functions may impact the replication mechanism. The Rep68/78 endonuclease activity or its phosphorylation may be differentially regulated depending on the helper virus. Further understanding of the replication mechanism may contribute to improve current systems for rAAV production, potentially increase rAAV yields and possibly inhibit undesired secondary responses in the host during treatment with rAAV vectors.

## Materials and Methods

### Cells and Viruses

Vero cells (African green monkey kidney) were obtained from ATCC (Manassas, Virginia, USA) and cultured in Dulbecco’s Modified Eagle Medium (DMEM) supplemented with 10% fetal bovine serum (FBS), 100 µg/ml streptomycin and 100 units/ml penicillin in a humified incubator at 37°C and 5% CO_2_. Vero 2-2 cells (71) were cultured in DMEM supplemented with 10% FBS, 100 µg/ml streptomycin, 100 units/ml penicillin and 500 µg/ml of G418 (G418 Sulfate, 10131035, Thermo Fisher Scientific, Waltham, MA, USA) in a humified incubator at 37°C and 5% CO_2_. Stocks of wt HSV-1 (strain F, provided by B. Rozman, University of Chicago) and HSVΔUL30 (HP66, provided by D. Coen, Harvard University, Boston, MA, USA) (72) were produced as previously described (73). Purified wt AAV2 was produced by the Viral Vector Facility (University of Zurich, Switzerland) as previously described (65,74) using the pAV2 plasmid (75). The recombinant AAV2 genome rAAVeCFPrep was constructed by replacing the mCherry coding sequence in the plasmid pAAVtCR (18) with the ECFP coding sequence from pECFP (Clontech, Mountain View, CA, USA). Purified recombinant AAV2 vectors, rAAVGFP, rAAVeCFPrep and rAAVmCherry (kindly provided by S. Sutter, University of Zurich, Switzerland), were produced by transient transfection of 293T cells with pDG (76) and pAAVeGFP (kindly provided by M. Linden, King’s College London School of Medicine, London, UK), pAAVeCFPrep or pAAVmCherry (kindly provided by J. Neidhardt, University of Zurich, Switzerland) and purified by an iodixanol density gradient. AdV5 was provided by U. Greber/M. Suomalainen (University of Zurich, Department of Molecular Life Sciences, Switzerland).

### Infection protocol

Cells were seeded the day before at 2*10^6^ cells per 10 cm tissue culture plate. For infection, cell culture medium was replaced with virus inoculum (viruses and corresponding multiplicity of infection (MOI) are indicated in the figure legends). After allowing the virus to adsorb for 30 min at 4°C, the cells were placed in a humified incubator for 1 h at 37°C and 5% CO_2_. Then, the virus inoculum was replaced with DMEM containing 2% FBS, and the cells were further incubated for the timepoints indicated in each experiment.

For combined infection and transfection experiments, the cells were infected as described above and, after the 1 h adsorption period at 37°C, transfected using Lipofectamine 2000 (11668019, Thermo Fisher Scientific, Waltham, MA, USA) with 10 µg of pCM-pol encoding HSV-1 UL30 or 10 µg of pCM-pol and 10 µg of pCM-UL42 encoding HSV-1 UL42 (Heilbronn, 1989, JVI) per plate according to the protocol provided by the supplier. At 24 h post infection (pi), FBS was added to reach a concentration of 2% in the medium, and the cells were further incubated at 37°C and 5% CO_2_ for 24 hpi.

### Southern Blot

Extrachromosomal DNA was extracted from infected cells using the Hirt protocol (39) and either digested extensively with *Hin*dIII, *Sal*I, or mung bean nuclease (MBN), or left untreated. The DNA was separated on 0.8% agarose gels and transferred onto nylon membranes (Hybond-N+, RPN203B, Amersham, Little Chalfont, UK). As size reference, DIG-labeled marker DNA was used (DNA molecular weight marker III, 11218602910, Roche). Hybridization was performed with probes specific for AAV2 *rep*, AAV2 ITR, ECFP/EGFP, or mCherry sequences. Detection with anti-digoxigenin antibody conjugated with alkaline phosphatase (Anti-Digoxigenin-AP Fab fragments, 11093274910, Roche, Switzerland) and activation with the chemiluminescent substrate CDP-Star (11759051001, Roche, Switzerland) was performed according to the manufacturers’ protocols. The *rep*, CFP/GFP and mCherry probes were synthesized using the PCR digoxigenin probe synthesis kit (1163090910, Roche, Switzerland) with primers shown in Table 1 and the following PCR conditions: 95°C for 2 min, followed by 95°C for 30 s, 55°C (ECFP/EGFP and mCherry) or 60°C (*rep*) for 30 s and 72°C for 40s for 30 cycles. The AAV2 ITR specific probe was labeled using the DIG DNA labeling kit (11175033910, Roche, Switzerland) using the following oligonucleotide: 5’-ttt ggt cgc ccg gcc tca gtg agc gag cga gcg cgc aga gag gga gtg gcc aa-3’. Chemiluminescence was visualized with the LI-COR imaging system Odyssey Fc (LI-COR Biosciences, NE, USA).

### Sequencing

Extrachromosomal DNA was extracted from infected cells using the Hirt protocol (39) and prepared for Nanopore sequencing using the ligation sequencing kit (LSK109, Oxford Nanopore Technologies, Oxford, UK) and the native barcoding expansion kit (EXP-NBD104, Oxford Nanopore Technologies, Oxford, UK) according to the manufacturers’ protocols. The samples were sequenced using a MinION sequencing device and flow cell (FLO-MIN_106, Oxford Nanopore Technologies, Oxford, UK). For PacBio sequencing, the Hirt DNA was first sheared and size selected for fragments >5 kb using the BluePippin device (Sage Science, MA, USA). Then libraries were prepared (100-938-900, SMRTbell® express template prep kit 2.0, PacBio, CA, USA) and sequenced on the PacBio Sequel II (PacBio, CA, USA) at the Functional Genomics Center (FGCZ, Zurich, Switzerland).

### Data analysis

Data analysis was performed as previously described in (Meier et al. 2021). Briefly, fastq-format files were transformed to fasta-format and subjected to blastn analysis against the AAV2 (Genbank #NC_001401) or rAAV genome sequences. Hits with e-values >0.1 were removed and the sum of the remaining hits were calculated for every read. Dot plots were drawn for reads with a sum of hits >3000. The dot plots were categorized manually. The code for the bioinformatics analysis can be found in Meier et al. (2021).

## Data availability

The raw sequencing data has been deposited at the National Center for Biotechnology Information Sequence Read Archive (NCBI SRA) website (https://www.ncbi.nlm.nih.gov/sra).

## Acknowledgments

We thank the members of the Fraefel lab for productive discussions. We thank Sereina Sutter for providing the Southern blot probes and rAAV. We thank Urs Greber and Maarit Suomalainen for providing AdV5.

---

**Fig. S1.**
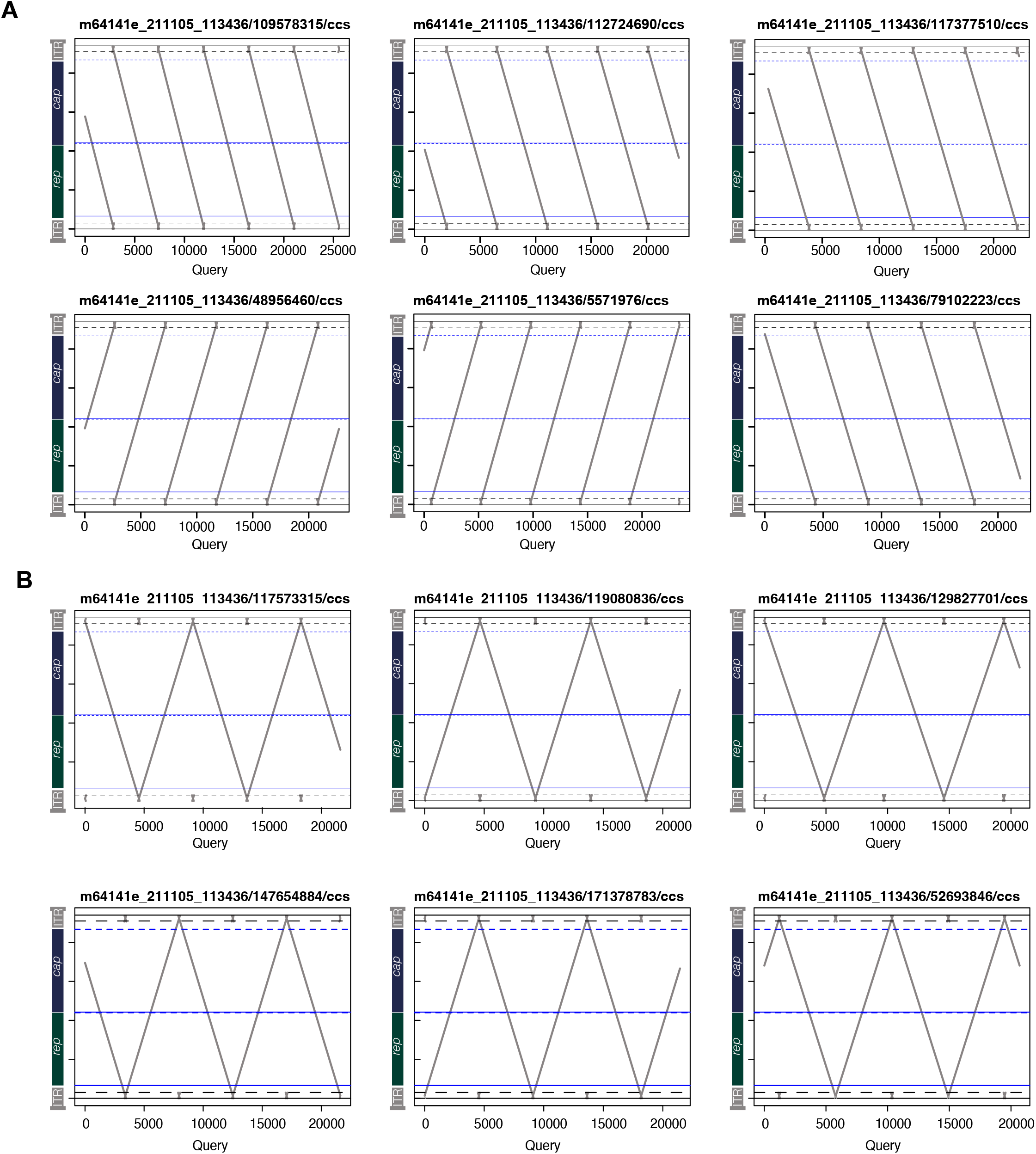
Dot plots of replication intermediates used for MSA analysis. Dot plots of the six (A) HT and (B) alternating reads used for MSA analysis.

**Fig. S2.**
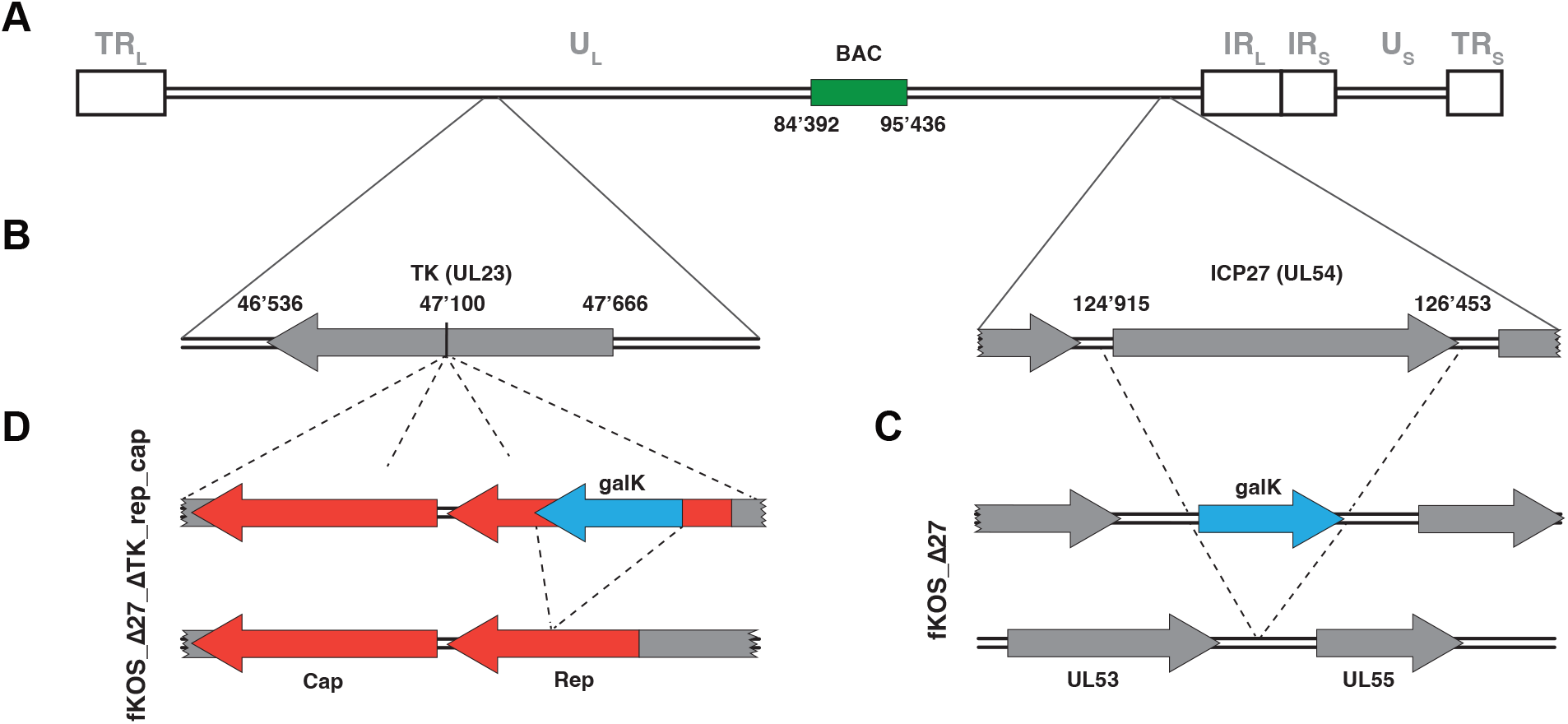
Schematic representation of the HSV KOS-37 BAC manipulations by recombineering. (A) Linear scheme of the circular KOS-37 BAC. (B) Magnified regions targeted by manipulations. (C) For deletion of ICP27, the galK expression cassette from pGalK (77) was amplified by PCR with the primers for_galK_d27 (5’-tgg cgc ttc act acg agc agg aga tcc aga ggc gcc tgt ttg atg tat gac ctg ttg aca atta at cat cgg ca-3’) and rev_galK_d27 (5’-aag gac aac acg tgg ggc gat ttg ttt gaa atg ttt tgt ttt tat tgt act cag cac tgt cct gct cct t-3’), and electroporated into *E.coli* SW102 harboring the HSV-1 KOS bacterial artificial chromosome (BAC). Electroporated bateria were selected on M9 minimal plates containing chloramphenicol (CAM) and galactose. Bacteria showing successful recombineering were induced and prepared as electrocompetent cell. Electroporation with double stranded (ds) 100bp oligos (ICP27_flank: 5’-tgg cgc ttg act acg agc agg aga tcc aga ggc gcc tgt ttg atg tat gag tac aat aaa aac aaa aca ttt caa aca aat cgc ccc acg tgt tgt cct t-3’) were introduced into the bacteria and selected on M9 minimal plates containing CAM and 2-deoxy-galactose (DOG). The resulting BAC was designated fKOS_Δ27. (D) Following ICP27 deletion and to insert the rep_cap9 cassette the galK expression cassette was amplified by PCR using the primers using the primers for_Hind3_galK (5’-tat aag ctt cct gtt gac aat taa tca tcg gca-3’), and rev_Hind3_galK (5’—gtg aag ctt cag cac tgt cct gct cct t-3’) with pGalK (77) as template. The amplimer was cut on both sides with *Hin*dIII and cloned into a plasmid which contains the *rep* and *cap* sequences of AAV9 and was synthesized at ATUM (Newark, CA). The part of the plasmid including the inserted galK expression cassette was excised from the plasmid, leaving TK-flanking regions on both sides of the rep_cap9 cassette, and inserted into *E.coli* SW102 cells carrying fKos_Δ27 and selected for galK positive colonies as described above. Finally, the galK cassett, situated within the AAV rep_cap9 cassette was deleted by recombineering with a ds 100 bp oligo (5’-caa acg ggt gcg cga gtc agt tgc gca gcc atc gac gtc aga cgc gga agc ttc gat ca acta cgc aga cag gta cca aaa caa atg ttc tcg tca cgt g-3’). The integrity of the manipulated region was confirmed by PCR amplicifaction and sequencing. The resulting BAC was designated fKOS Δ27 ΔTK rep/cap. To reconstitute recombinant HSV isolated HSV-1 BAC DNA fKOSΔ27ΔTK rep/cap was treated with Cre-recombinase (M0298SNEB, Ipswich, MA, USA) for 30 min at 30°C. Subsequently, the BAC DNA was transfected into Vero 2-2 cells. Progeny virus was harvested and used for infection of fresh cells. The Cre-mediated recombination was confirmed by visualization of absent GFP expression. The resulting recombinant HSV-1 vectors were designated rHSV-1 rep/cap (HSV-1RC).

**Fig. S3.**
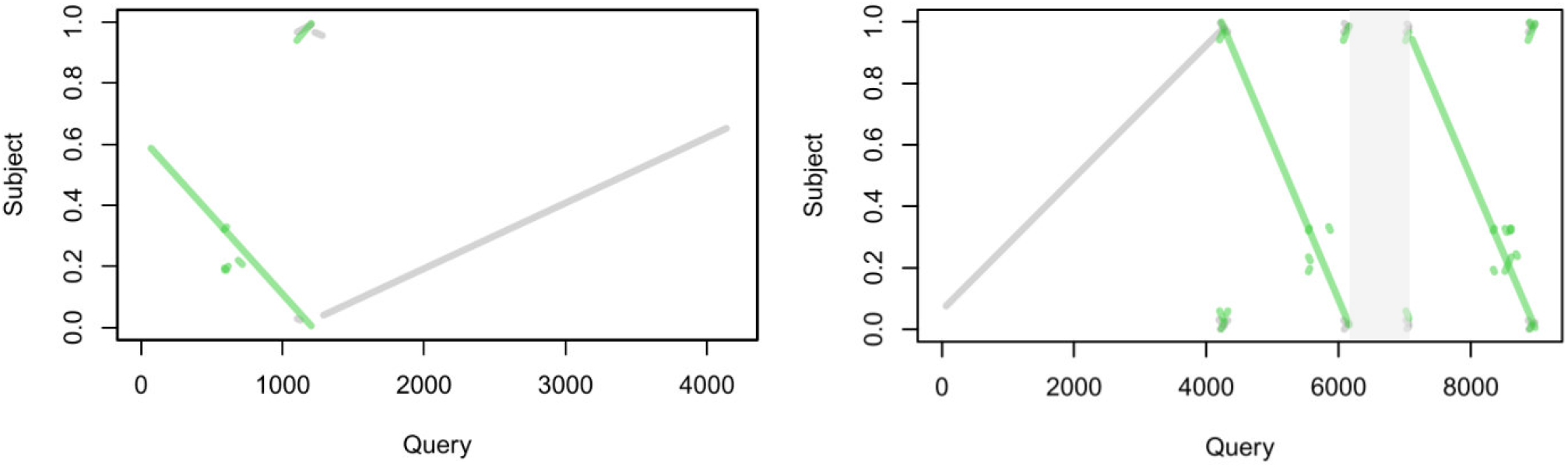
Southern blot and nanopore read analysis of recombination efficiency. Sequencing read analysis of Hirt DNA from cells co-infected with AAV2, rAAVGFP and AdV5. Two dotblots from two reads of a total of 6399 reads that show recombination are illustrated. Recombination ratio amounts to 0.03%. The sequence indicated in grey is of unknown origin.

## References

1. Samulski RJ, Muzyczka N. AAV-Mediated Gene Therapy for Research and Therapeutic Purposes. Annu Rev Virol. 2014 Nov 3;1(1):427–51.

2. Lusby E, Fife KH, Berns KI. Nucleotide sequence of the inverted terminal repetition in adeno-associated virus DNA. J Virol [Internet]. 1980;34(2):402–9. Available from: https://pubmed.ncbi.nlm.nih.gov/6246271/

3. Xiao X, Xiao W, Li J, Samulski RJ. A novel 165-base-pair terminal repeat sequence is the sole cis requirement for the adeno-associated virus life cycle. J Virol. 1997;71(2):941–8.

4. Samulski RJ, Berns KI, Tan M, Muzyczka N. Cloning of adeno-associated virus into pBR322: Rescue of intact virus from the recombinant plasmid in human cells. Proc Natl Acad Sci U S A [Internet]. 1982;79(6 I):2077–81. Available from: https://pubmed.ncbi.nlm.nih.gov/6281795/

5. Samulski RJ, Srivastava A, Berns KI, Muzyczka N. Rescue of adeno-associated virus from recombinant plasmids: gene correction within the terminal repeats of AAV. Cell [Internet]. 1983 [cited 2023 Mar 27];33(1):135–43. Available from: https://pubmed.ncbi.nlm.nih.gov/6088052/

6. Yakobson B, Hrynko TA, Peak MJ, Winocour E. Replication of adeno-associated virus in cells irradiated with UV light at 254 nm. J Virol. 1989;63(3):1023–30.

7. Yakobson B, Koch T, Winocour E. Replication of adeno-associated virus in synchronized cells without the addition of a helper virus. J Virol [Internet]. 1987;61(4):972–81. Available from: /pmc/articles/PMC254052/?report=abstract

8. Georg-Fries B, Biederlack S, Wolf J, Zur Hausen H. Analysis of proteins, helper dependence, and seroepidemiology of a new human parvovirus. Virology. 1984 Apr;134(1):64–71.

9. Buller RML, Janik JE, Sebring ED, Rose JA. Herpes simplex virus types 1 and 2 completely help adenovirus-associated virus replication. J Virol [Internet]. 1981 Oct;40(1):241–7. Available from: https://pubmed.ncbi.nlm.nih.gov/6270377/

10. Handa H, Shiroki K, Shimojo H. Helper factor(s) for growth of adeno-associated virus in cells transformed by adenovirus 12. Proc Natl Acad Sci U S A [Internet]. 1977;74(10):4508–10. Available from: https://www.pnas.org

11. Heilbronn R, Schlehofer JR, Yalklnoglu AO, Hausen H Zur. Selective dna-amplification induced by carcinogens (initiators): Evidence for a role of proteases and DNA polymerase alpha. Int J Cancer [Internet]. 1985;36(1):85–91. Available from: https://pubmed.ncbi.nlm.nih.gov/3894246/

12. Taylor TJ, Brockman MA, McNamee EE, Knipe DM. Herpes simplex virus. Front Biosci [Internet]. 2002 Mar;7(4):752–64. Available from: https://www.imrpress.com/journal/FBL/7/4/10.2741/taylor

13. Poffenberger KL, Roizman B. A noninverting genome of a viable herpes simplex virus 1: presence of head-to-tail linkages in packaged genomes and requirements for circularization after infection. J Virol [Internet]. 1985 Feb;53(2):587–95. Available from: https://pubmed.ncbi.nlm.nih.gov/2982037/

14. Vlazny DA, Frenkel N. Replication of herpes simplex virus DNA: localization of replication recognition signals within defective virus genomes. Proc Natl Acad Sci U S A [Internet]. 1981;78(2):742–6. Available from: https://pubmed.ncbi.nlm.nih.gov/6262768/

15. Weerasooriya S, DiScipio KA, Darwish AS, Bai P, Weller SK. Herpes simplex virus 1 ICP8 mutant lacking annealing activity is deficient for viral DNA replication. Proc Natl Acad Sci U S A [Internet]. 2019 Jan;116(3):1033–42. Available from: www.pnas.org/cgi/doi/10.1073/pnas.1817642116

16. Schumacher AJ, Mohni KN, Kan Y, Hendrickson EA, Stark JM, Weller SK. The HSV-1 Exonuclease, UL12, Stimulates Recombination by a Single Strand Annealing Mechanism. PLoS Pathog [Internet]. 2012 Aug [cited 2022 Sep 14];8(8):e1002862. Available from: https://journals.plos.org/plospathogens/article?id=10.1371/journal.ppat.1002862

17. Weindler FW, Heilbronn R. A subset of herpes simplex virus replication genes provides helper functions for productive adeno-associated virus replication. J Virol [Internet]. 1991;65(5):2476–83. Available from: https://pubmed.ncbi.nlm.nih.gov/1850024/

18. Alazard-Dany N, Nicolas A, Ploquin A, Strasser R, Greco A, Epstein AL, et al. Definition of Herpes Simplex Virus Type 1 Helper Activities for Adeno-Associated Virus Early Replication Events. PLoS Pathog [Internet]. 2009;5(3):e1000340. Available from: https://journals.plos.org/plospathogens/article?id=10.1371/journal.ppat.1000340

19. Handa H, Carter BJ. Adeno-associated virus DNA replication complexes in herpes simplex virus or adenovirus-infected cells. Journal of Biological Chemistry. 1979 Jul;254(14):6603–10.

20. Doerfler W, Böhm P. Adenoviruses: Model and Vectors in Virus-Host Interactions: Virion-Structure, Viral Replication and Host-Cell Interactions. 2003.

21. Hoeben RC, Uil TG. Adenovirus DNA Replication. Cold Spring Harb Perspect Biol [Internet]. 2013 Mar 1 [cited 2023 Sep 20];5(3):a013003. Available from: http://cshperspectives.cshlp.org/content/5/3/a013003.full

22. Laughlin CA, Jones N, Carter BJ. Effect of deletions in adenovirus early region 1 genes upon replication of adeno-associated virus. J Virol [Internet]. 1982 Mar;41(3):868–76. Available from: https://pubmed.ncbi.nlm.nih.gov/6284977/

23. Tratschin JD, Miller IL, Carter BJ. Genetic analysis of adeno-associated virus: properties of deletion mutants constructed in vitro and evidence for an adeno-associated virus replication function. J Virol [Internet]. 1984 Sep;51(3):611–9. Available from: https://pubmed.ncbi.nlm.nih.gov/6088786/

24. Chang LS, Shi Y, Shenk T. Adeno-associated virus P5 promoter contains an adenovirus E1A-inducible element and a binding site for the major late transcription factor. J Virol [Internet]. 1989 Aug;63(8):3479–88. Available from: https://pubmed.ncbi.nlm.nih.gov/2545917/

25. Samulski RJ, Shenk T. Adenovirus E1B 55-Mr polypeptide facilitates timely cytoplasmic accumulation of adeno-associated virus mRNAs. J Virol [Internet]. 1988 Jan;62(1):206–10. Available from: https://pubmed.ncbi.nlm.nih.gov/2824848/

26. Fisher KJ, Gao GP, Weitzman MD, DeMatteo R, Burda JF, Wilson JM. Transduction with recombinant adeno-associated virus for gene therapy is limited by leading-strand synthesis. J Virol [Internet]. 1996 Jan;70(1):520–32. Available from: https://pubmed.ncbi.nlm.nih.gov/8523565/

27. Janik JE, Huston MM, Cho K, Rose JA. Efficient synthesis of adeno-associated virus structural proteins requires both adenovirus DNA binding protein and VA I RNA. Virology [Internet]. 1989;168(2):320–9. Available from: https://pubmed.ncbi.nlm.nih.gov/2536986/

28. Nash K, Chen W, McDonald WF, Zhou X, Muzyczka N. Purification of Host Cell Enzymes Involved in Adeno-Associated Virus DNA Replication. J Virol [Internet]. 2007 Jun;81(11):5777. Available from: /pmc/articles/PMC1900299/

29. Nash K, Chen W, Muzyczka N. Complete In Vitro Reconstitution of Adeno-Associated Virus DNA Replication Requires the Minichromosome Maintenance Complex Proteins. J Virol [Internet]. 2008 Feb;82(3):1458–64. Available from: https://journals.asm.org/doi/10.1128/JVI.01968-07

30. Knipe David M. HPM. Fields of Virology, Volume I. Vol. 1. 2013. 2582 p.

31. Im DS, Muzyczka N. The AAV origin binding protein Rep68 is an ATP-dependent site-specific endonuclease with DNA helicase activity. Cell [Internet]. 1990 May;61(3):447–57. Available from: https://pubmed.ncbi.nlm.nih.gov/2159383/

32. King JA, Dubielzig R, Grimm D, Kleinschmidt JA. DNA helicase-mediated packaging of adeno-associated virus type 2 genomes into preformed capsids. EMBO Journal [Internet]. 2001 Jun;20(12):3282–91. Available from: /pmc/articles/PMC150213/?report=abstract

33. Lusby E, Bohenzky R, Berns KI. Inverted terminal repetition in adeno-associated virus DNA: independence of the orientation at either end of the genome. J Virol [Internet]. 1981 Mar [cited 2023 Jul 18];37(3):1083–6. Available from: https://pubmed.ncbi.nlm.nih.gov/6262528/

34. Hauswirth WW, Berns KI. Origin and termination of adeno-associated virus DNA replication. Virology [Internet]. 1977 [cited 2023 Sep 20];78(2):488–99. Available from: https://pubmed.ncbi.nlm.nih.gov/867815/

35. Straus SE, Sebring ED, Rose JA. Concatemers of alternating plus and minus strands are intermediates in adenovirus-associated virus DNA synthesis. Proc Natl Acad Sci U S A [Internet]. 1976 [cited 2023 Jul 5];73(3):742–6. Available from: https://pubmed.ncbi.nlm.nih.gov/1062784/

36. Snyder RO, Samulski RJ, Muzyczka N. In vitro resolution of covalently joined AAV chromosome ends. Cell [Internet]. 1990 Jan;60(1):105–13. Available from: https://pubmed.ncbi.nlm.nih.gov/2153052/

37. Brister JR, Muzyczka N. Mechanism of Rep-mediated adeno-associated virus origin nicking. J Virol [Internet]. 2000 Sep;74(17):7762–71. Available from: https://pubmed.ncbi.nlm.nih.gov/10933682/

38. Meier AF, Tobler K, Leisi R, Lkharrazi A, Ros C, Fraefel C. Herpes simplex virus co-infection facilitates rolling circle replication of the adeno-associated virus genome. PLoS Pathog. 2021;17(6):e1009638.

39. Hirt B. Selective extraction of polyoma DNA from infected mouse cell cultures. J Mol Biol. 1967 Jun 14;26(2):365–9.

40. Wang Y, Zhao Y, Bollas A, Wang Y, Au KF. Nanopore sequencing technology, bioinformatics and applications. Nat Biotechnol [Internet]. 2021 Nov 1 [cited 2023 May 25];39(11):1348–65. Available from: https://pubmed.ncbi.nlm.nih.gov/34750572/

41. Rhoads A, Au KF. PacBio Sequencing and Its Applications. Genomics Proteomics Bioinformatics [Internet]. 2015 [cited 2023 May 25];13(5):278–89. Available from: https://pubmed.ncbi.nlm.nih.gov/26542840/

42. Nash K, Chen W, Salganik M, Muzyczka N. Identification of Cellular Proteins That Interact with the Adeno-Associated Virus Rep Protein. J Virol [Internet]. 2009 Jan [cited 2023 May 12];83(1):454–69. Available from: https://journals.asm.org/doi/10.1128/JVI.01939-08

43. Nicolas A, Alazard-Dany N, Biollay C, Arata L, Jolinon N, Kuhn L, et al. Identification of Rep-Associated Factors in Herpes Simplex Virus Type 1-Induced Adeno-Associated Virus Type 2 Replication Compartments. J Virol [Internet]. 2010 Sep [cited 2023 May 3];84(17):8871–87. Available from: https://journals.asm.org/doi/10.1128/JVI.00725-10

44. Ward P, Falkenberg M, Elias P, Weitzman M, Linden RM. Rep-Dependent Initiation of Adeno-Associated Virus Type 2 DNA Replication by a Herpes Simplex Virus Type 1 Replication Complex in a Reconstituted System. J Virol [Internet]. 2001 Nov [cited 2023 Mar 27];75(21):10250. Available from: /pmc/articles/PMC114599/

45. Tattersall P, Ward DC. Rolling hairpin model for replication of parvovirus and linear chromosomal DNA. Nature [Internet]. 1976 [cited 2023 Jul 5];263(5573):106–9. Available from: https://pubmed.ncbi.nlm.nih.gov/967244/

46. Berns KI. Parvovirus replication. Microbiol Rev [Internet]. 1990 Sep [cited 2023 Jul 5];54(3):316–29. Available from: https://pubmed.ncbi.nlm.nih.gov/2215424/

47. Ward P, Dean FB, O’Donnell ME, Berns KI. Role of the adenovirus DNA-binding protein in in vitro adeno-associated virus DNA replication. J Virol [Internet]. 1998 Jan [cited 2023 Jul 12];72(1):420–7. Available from: https://pubmed.ncbi.nlm.nih.gov/9420241/

48. Xiao X, Li J, Samulski RJ. Production of high-titer recombinant adeno-associated virus vectors in the absence of helper adenovirus. J Virol [Internet]. 1998 Mar [cited 2023 Jul 10];72(3):2224–32. Available from: https://pubmed.ncbi.nlm.nih.gov/9499080/

49. Skaliter R, Lehman IR. Rolling circle DNA replication in vitro by a complex of herpes simplex virus type 1-encoded enzymes. Proc Natl Acad Sci U S A [Internet]. 1994 Oct 25 [cited 2023 May 31];91(22):10665–9. Available from: https://pubmed.ncbi.nlm.nih.gov/7938010/

50. Choi VW, Samulski RJ, McCarty DM. Effects of Adeno-Associated Virus DNA Hairpin Structure on Recombination. J Virol. 2005 Jun;79(11):6801–7.

51. Duan D, Yan Z, Yue Y, Engelhardt JF. Structural analysis of adeno-associated virus transduction circular intermediates. Virology. 1999 Aug;261(1):8–14.

52. Sun X, Lu Y, Bish LT, Calcedo R, Wilson JM, Gao G. Molecular analysis of vector genome structures after liver transduction by conventional and self-complementary adeno-associated viral serotype vectors in murine and nonhuman primate models. Hum Gene Ther [Internet]. 2010 Jun 1 [cited 2023 Jun 7];21(6):750–62. Available from: https://pubmed.ncbi.nlm.nih.gov/20113166/

53. Weiden MD, Ginsberg HS. Deletion of the E4 region of the genome produces adenovirus DNA concatemers. Proc Natl Acad Sci U S A [Internet]. 1994 Jan;91(1):153–7. Available from: https://pubmed.ncbi.nlm.nih.gov/8278357/

54. Stracker TH, Carson CT, Weilzman MD. Adenovirus oncoproteins inactivate the Mre11-Rad50-NBS1 DNA repair complex. Nature [Internet]. 2002 Jul;418(6895):348–52. Available from: https://pubmed.ncbi.nlm.nih.gov/12124628/

55. Allen JM, Halbert CL, Miller AD. Improved adeno-associated virus vector production with transfection of a single helper adenovirus gene, E4orf6. Mol Ther [Internet]. 2000 Jan [cited 2023 Jul 10];1(1):88–95. Available from: https://pubmed.ncbi.nlm.nih.gov/10933916/

56. Musatov S, Roberts J, Pfaff D, Kaplitt M. A cis-Acting Element That Directs Circular Adeno-Associated Virus Replication and Packaging. J Virol [Internet]. 2002 Dec 15 [cited 2022 Sep 14];76(24):12792. Available from: /pmc/articles/PMC136660/

57. Hirsch ML, Fagan BM, Dumitru R, Bower JJ, Yadav S, Porteus MH, et al. Viral Single-Strand DNA Induces p53-Dependent Apoptosis in Human Embryonic Stem Cells. PLoS One [Internet]. 2011 [cited 2023 Jul 12];6(11):27520. Available from: /pmc/articles/PMC3219675/

58. Raj K, Ogston P, Beard P. Virus-mediated killing of cells that lack p53 activity. Nature 2001 412:6850 [Internet]. 2001 Aug 30 [cited 2023 Aug 14];412(6850):914–7. Available from: https://www.nature.com/articles/35091082

59. Bucher K, Rodríguez-Bocanegra E, Wissinger B, Strasser T, Clark SJ, Birkenfeld AL, et al. Extra-viral DNA in adeno-associated viral vector preparations induces TLR9-dependent innate immune responses in human plasmacytoid dendritic cells. Sci Rep. 2023;13(1):1890.

60. Inagaki K, Ma C, Storm TA, Kay MA, Nakai H. The Role of DNA-PKcs and Artemis in Opening Viral DNA Hairpin Termini in Various Tissues in Mice. J Virol [Internet]. 2007 Oct 15 [cited 2023 Jul 5];81(20):11304–21. Available from: https://journals.asm.org/doi/10.1128/jvi.01225-07

61. Zhang X, Succi J, Feng Z, Prithivirajsingh S, Story MD, Legerski RJ. Artemis Is a Phosphorylation Target of ATM and ATR and Is Involved in the G2/M DNA Damage Checkpoint Response. Mol Cell Biol [Internet]. 2004 Oct 1 [cited 2023 Nov 3];24(20):9207. Available from: /pmc/articles/PMC517881/

62. Schwartz RA, Carson CT, Schuberth C, Weitzman MD. Adeno-Associated Virus Replication Induces a DNA Damage Response Coordinated by DNA-Dependent Protein Kinase. J Virol. 2009;83(12):6269.

63. Parkinson J, Lees-Miller SP, Everett RD. Herpes simplex virus type 1 immediate-early protein vmw110 induces the proteasome-dependent degradation of the catalytic subunit of DNA-dependent protein kinase. J Virol [Internet]. 1999 Jan [cited 2023 Jul 5];73(1):650–7. Available from: https://pubmed.ncbi.nlm.nih.gov/9847370/

64. Lees-Miller SP, Long MC, Kilvert MA, Lam V, Rice SA, Spencer CA. Attenuation of DNA-dependent protein kinase activity and its catalytic subunit by the herpes simplex virus type 1 transactivator ICP0. J Virol [Internet]. 1996 Nov [cited 2023 Jul 5];70(11):7471–7. Available from: https://pubmed.ncbi.nlm.nih.gov/8892865/

65. Vogel R, Seyffert M, Strasser R, de Oliveira AP, Dresch C, Glauser DL, et al. Adeno-Associated Virus Type 2 Modulates the Host DNA Damage Response Induced by Herpes Simplex Virus 1 during Coinfection. J Virol [Internet]. 2012 Jan;86(1):143–55. Available from: http://jvi.asm.org/.

66. Gilbert W, Dressler D. DNA Replication: The Rolling Circle Model. Cold Spring Harb Symp Quant Biol [Internet]. 1968 Jan 1 [cited 2023 Jul 7];33:473–84. Available from: http://symposium.cshlp.org/content/33/473

67. Song S, Laipis PJ, Berns KI, Flotte TR. Effect of DNA-dependent protein kinase on the molecular fate of the rAAV2 genome in skeletal muscle. Proc Natl Acad Sci U S A [Internet]. 2001 Mar 27 [cited 2023 Jul 7];98(7):4084–8. Available from: https://www.pnas.org/doi/abs/10.1073/pnas.061014598

68. Nakai H, Storm TA, Fuess S, Kay MA. Pathways of removal of free DNA vector ends in normal and DNA-PKcs-deficient SCID mouse hepatocytes transduced with rAAV vectors. Hum Gene Ther [Internet]. 2003 Jun 10 [cited 2023 Jul 7];14(9):871–81. Available from: https://pubmed.ncbi.nlm.nih.gov/12828858/

69. Duan D, Yue Y, Engelhardt JF. Consequences of DNA-Dependent Protein Kinase Catalytic Subunit Deficiency on Recombinant Adeno-Associated Virus Genome Circularization and Heterodimerization in Muscle Tissue. J Virol [Internet]. 2003 Apr 15 [cited 2023 Jul 7];77(8):4751–9. Available from: https://journals.asm.org/doi/10.1128/jvi.77.8.4751-4759.2003

70. Fragkos M, Breuleux M, Clément N, Beard P. Recombinant Adeno-Associated Viral Vectors Are Deficient in Provoking a DNA Damage Response. J Virol. 2008;82(15):7379–87.

71. Smith IL, Hardwicke MA, Sandri-Goldin RM. Evidence that the herpes simplex virus immediate early protein ICP27 acts post-transcriptionally during infection to regulate gene expression. Virology [Internet]. 1992 [cited 2023 May 26];186(1):74–86. Available from: https://pubmed.ncbi.nlm.nih.gov/1309283/

72. Marcy AI, Yager DR, Coen DM. Isolation and characterization of herpes simplex virus mutants containing engineered mutations at the DNA polymerase locus. J Virol [Internet]. 1990 May;64(5):2208–16. Available from: https://journals.asm.org/journal/jvi

73. Sutter SO, Marconi P, Meier AF. Herpes Simplex Virus Growth, Preparation, and Assay. Methods in Molecular Biology [Internet]. 2020;2060:57–72. Available from: https://link.springer.com/protocol/10.1007/978-1-4939-9814-2_3

74. Zolotukhin S, Byrne BJ, Mason E, Zolotukhin I, Potter M, Chesnut K, et al. Recombinant adeno-associated virus purification using novel methods improves infectious titer and yield. Gene Ther [Internet]. 1999 Jun [cited 2023 May 26];6(6):973–85. Available from: https://pubmed.ncbi.nlm.nih.gov/10455399/

75. Laughlin CA, Tratschin JD, Coon H, Carter BJ. Cloning of infectious adeno-associated virus genomes in bacterial plasmids. Gene. 1983 Jul 1;23(1):65–73.

76. Grimm D, Kern A, Rittner K, Kleinschmidt JA. Novel tools for production and purification of recombinant adenoassociated virus vectors. Hum Gene Ther [Internet]. 1998 Dec 10 [cited 2023 Jun 16];9(18):2745–60. Available from: https://pubmed.ncbi.nlm.nih.gov/9874273/

